# The First Experience of Accelerator-based Lithium Neutron Capture Therapy *in vivo*

**DOI:** 10.64898/2026.09.04.749316

**Authors:** Iuliia Taskaeva, Anna Kasatova, Timofey Bykov, Vladimir Richter, Evgeniia Sokolova, Nataliya Bgatova, Sergey Taskaev

## Abstract

Boron neutron capture therapy (BNCT) is one of the promising treatment methods for cancer. BNCT is based on the unique high ability of the non-radioactive boron-10 (^10^B) nucleus to absorb thermal neutrons. The absorption of a neutron by boron results in a nuclear reaction ^10^B(n,α)^7^Li with 84% of the energy being released within the cell, thereby inducing cancer cell death. We propose a novel approach based on using lithium instead of boron in neutron capture therapy: the neutron capture reaction cross-section for lithium is 4 times smaller than that of boron, while the energy release is 2 times higher, and the primary advantage of lithium is that 100% of the energy is released inside the cell. A series of in vivo neutron irradiation experiments using the murine B16 skin melanoma model demonstrated the efficacy of lithium neutron capture therapy (LiNCT). Higher lithium concentrations in the tumor achieved through intraperitoneal administration, compared to the oral route, contributed to a significant increase in animal survival and a significant decrease in tumor growth. The results of our study provide entirely new opportunities for the development of neutron capture therapy, both with lithium and with a possible lithium-boron combination.

## Introduction

Boron neutron capture therapy (BNCT) is a radiotherapeutic approach based on the nuclear reaction ^10^B(n,α)^7^Li, in which thermal neutron absorption by boron-10 (^10^B) generates high-linear energy transfer (LET) particles – an alpha particle (α) and a lithium-7 (^7^Li) nucleus – depositing cytotoxic energy within a limited range (∼10 μm)^1,2^. This selective mechanism enables localized tumor cell destruction while sparing adjacent healthy tissues^3^.

The conception of neutron capture therapy (NCT) was first proposed by Locher^4^, yet clinical implementation became feasible only after the end of the Second World War using the advent of nuclear reactors^5^. Despite promising clinical outcomes, BNCT faces several limitations hindering its widespread adoption^6,7^. A critical challenge is achieving sufficient and selective ^10^B accumulation in tumor cells while minimizing off-target retention in blood and normal tissues^6^. Currently, although novel boron-delivery agents are under investigation, the clinical protocols rely primarily on second-generation compounds, such as boronophenylalanine (BPA) and sodium borocaptate (BSH)^8,9^.

The use of lithium instead of boron in NCT represents a promising therapeutic approach due to the physical properties of lithium. The lithium-6 isotope (^6^Li) has a large thermal neutron absorption cross-section (940 barns)^10^, and the nuclear reaction (^6^Li(n,α)^3^H) produces LET particles, which release energy in a localized volume (∼10 μm). In contrast, other isotopes with high absorption cross-sections (e.g., ^113^Cd, ^135^Xe, ^149^Sm, ^151^Eu, ^155^Gd, ^157^Gd, ^174^Hf, ^199^Hg) preferentially undergo (n,γ)-reactions, which lack a comparable local energy release. Moreover, the neutron capture reaction cross-section for lithium is 4 times smaller than that of boron, while the energy release is 2 times higher, and the main advantage of lithium is that 100% of the energy will be released inside the cell.

After John Cade demonstrated the efficacy of lithium therapy for mania in 1949, lithium salts began to be actively researched and used in medicine^11^. Despite initial concerns regarding its narrow therapeutic index, subsequent pharmacokinetic studies and standardized monitoring protocols ensured its safe clinical use, leading to US Food and Drug Administration agency approval in the 1970s^12^. Today, lithium salts (e.g., carbonate, chloride, citrate) remain first-line treatments for bipolar disorder^13,14^. Although lithium’s toxicity profile requires careful dose management, advances in pharmacokinetics and therapeutic drug monitoring have significantly reduced adverse effects^15,16^. These well-established pharmacological protocols may facilitate the introducing of lithium to NCT by providing precise control of tissue lithium concentrations.

The concept of lithium using for NCT can be traced back to Locher’s proposition in 1936: “*In particular, there exist the possibilities of introducing small quantities of strong neutron absorbers into the regions where it is desired to liberate ionization energy (a simple illustration would be the injection of a soluble, non-toxic compound of boron, lithium, gadolinium, or gold into a superficial cancer, followed by bombardment with slow neutrons)*”^4^. To date, the number of publications regarding LiNCT remains very limited. The first attempt to perform LiNCT *in vivo* was conducted by Zahl and co-authors, in which murine sarcomas were irradiated with slow neutrons, resulting in substantial tumor regression^17^. Subsequently, it was demonstrated that lithium is efficiently accumulated by glioma and neuroblastoma cells *in vitro*^18,19^.

A renewed interest in LiNCT was sparked by Rendina in 2010. In his publication, Rendina reasonably noted that the development of boron-containing compound chemistry, optimization of neutron beams, and the higher neutron capture cross-section of boron compared to lithium have directed the evolution of NCT towards boron over several decades, with lithium largely overlooked^20^. However, these advancements did not lead to considerable outcomes regarding lithium-based approaches. In 2023, Gonçalves and colleagues published an elegant study in which they developed ^6^Li-filled carbon nanohorns and demonstrated their effectiveness for *in vitro* LiNCT using rat osteosarcoma cells^21^. Finally, our previous studies showed effective accumulation of lithium in melanoma cells following the administration of various lithium salts containing the natural lithium. These studies also clearly demonstrated the absence of toxicity after a single administration of high-dose lithium for the of LiNCT^22-24^. In the current study, we aimed to assess the anticancer efficacy of LiNCT using a mouse melanoma model. To the best of our knowledge, this is the first report to assess the LiNCT efficacy *in vivo* using accelerator-based neutron beam suitable for clinical implementation of neutron capture therapy.

## Results

### Lithium biodistribution and LiNCT results after peroral lithium administration

Lithium concentrations in tumor, blood, muscle, skin, liver, kidney and brain after peroral lithium administration was measured using the ICP-AES method (table 1). The highest accumulation of lithium in the tumor was achieved 4 hours after oral administration of lithium chloride (30.2 µg/g). At this time point, the tumor/skin lithium maximal concentration ratio (T/S) was 4.1, and the tumor/blood ratio (T/B) reached 1.5. High lithium content in the tumor was also detected at the 1- hour time point, however, the achieved tumor/skin and tumor/blood ratios were lower than those at the 4-hour time point. It should be noted that lithium concentrations in other organs were lower than in the tumor 4 hours after oral administration of lithium chloride, specifically, lithium accumulation in kidneys was 27.4 μg/g, and in liver – 9.4 μg/g. Kidneys maintained consistently high lithium levels (24-36 μg/g) during the entire monitoring window, whereas brain concentrations never surpassed 6.2 μg/g which correlates with data from our previous lithium biodistribution data.

**Table 1.**
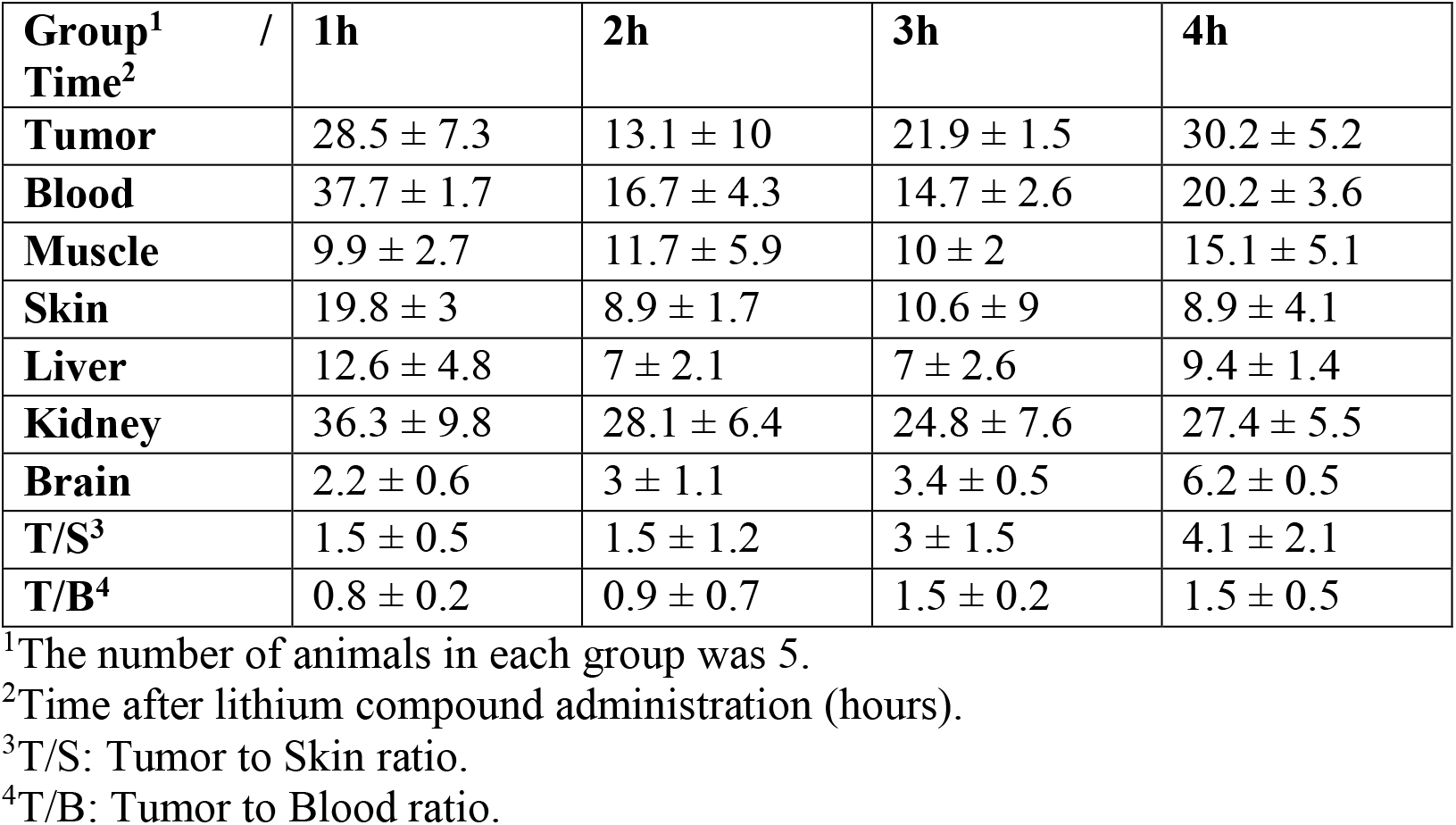
The lithium concentrations in tissues of the B16 melanoma-bearing mice after peroral lithium chloride administration (M±SD)

| **Group<sup>1</sup>**<br>**Time<sup>2</sup>** / | **1h** | **2h** | **3h** | **4h** |
| --- | --- | --- | --- | --- |
| **Tumor** | 28.5 ± 7.3 | 13.1 ± 10 | 21.9 ± 1.5 | 30.2 ± 5.2 |
| **Blood** | 37.7 ± 1.7 | 16.7 ± 4.3 | 14.7 ± 2.6 | 20.2 ± 3.6 |
| **Muscle** | 9.9 ± 2.7 | 11.7 ± 5.9 | 10 ± 2 | 15.1 ± 5.1 |
| **Skin** | 19.8 ± 3 | 8.9 ± 1.7 | 10.6 ± 9 | 8.9 ± 4.1 |
| **Liver** | 12.6 ± 4.8 | 7 ± 2.1 | 7 ± 2.6 | 9.4 ± 1.4 |
| **Kidney** | 36.3 ± 9.8 | 28.1 ± 6.4 | 24.8 ± 7.6 | 27.4 ± 5.5 |
| **Brain** | 2.2 ± 0.6 | 3 ± 1.1 | 3.4 ± 0.5 | 6.2 ± 0.5 |
| **T/S<sup>3</sup>** | 1.5 ± 0.5 | 1.5 ± 1.2 | 3 ± 1.5 | 4.1 ± 2.1 |
| **T/B<sup>4</sup>** | 0.8 ± 0.2 | 0.9 ± 0.7 | 1.5 ± 0.2 | 1.5 ± 0.5 |

<sup>1</sup>The number of animals in each group was 5.
<sup>2</sup>Time after lithium compound administration (hours).
<sup>3</sup>T/S: Tumor to Skin ratio.
<sup>4</sup>T/B: Tumor to Blood ratio.

Figure 1 shows the animal weights, Kaplan-Meier survival curves, and tumor growth dynamics for each group after lithium administration *per os*. The weight of mice irradiated with neutrons was lower than in the Control and Li groups, which could be due to the significantly larger tumor volume in these animals. The log-rank test indicated no statistically significant difference in survival among the groups (p = 0.086), although a trend towards improved survival in the Neu and LiNCT groups was observed. To analyze the tumor growth dynamics, GLMM analysis was performed taking into account the interaction of group and time; a trend toward decreased tumor growth was also observed in the LiNCT group in the first 3 weeks of the experiment, but no significant differences were found compared to the control groups (Control (p = 0.093), Li (p = 0.928), Neu (p = 0.837)).

**Figure 1.**
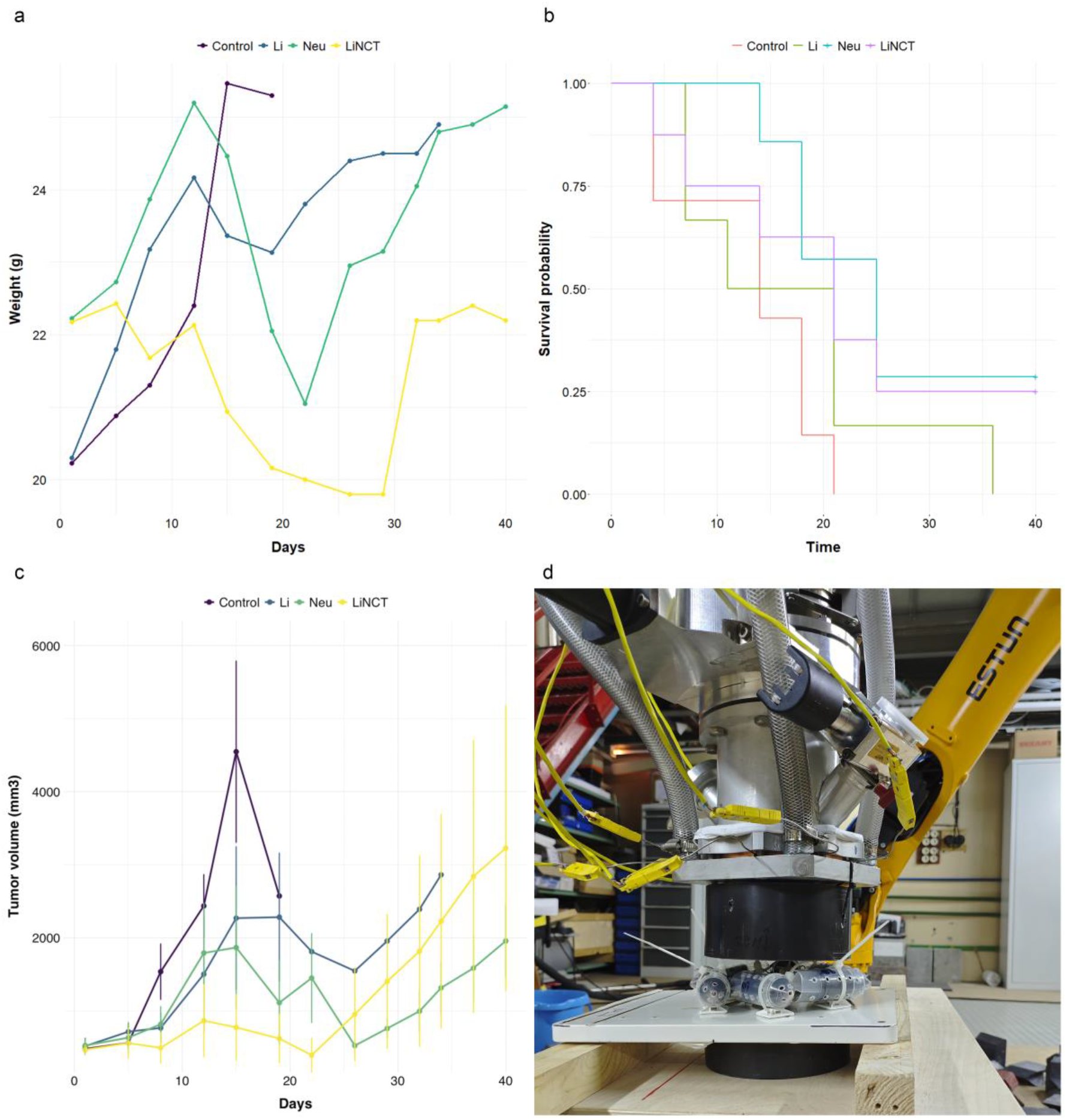
Evaluation of survival rate, animal weight and tumor growth dynamics for each experimental group after peroral lithium administration. (**a**) This figure shows the results of the animal weight dynamics (mean values) for B16 skin melanoma model after LiNCT and in control groups. (**b**) Survival times (days) are plotted, statistical analysis was performed using the log-rank test with Holm’s correction for multiple comparisons (“Control”: n = 7, “Li”: n = 6, “Neu”: n = 7, “LiNCT”: n = 8). (**c**) Tumor volume dynamics was analyzed using a generalized linear mixedeffects model (GLMM), the data are presented as means ± SD. (**d**) The representative image of experimental animals positioned in a specially designed restrictor under a collimator for neutron irradiation.

### Lithium biodistribution and LiNCT efficiency after intraperitoneal lithium administration

Lithium biodistribution data after intraperitoneal injection of lithium chloride is presented in table 2. The highest lithium concentrations in the tumor were observed one hour after lithium chloride injection and measured 54.1 µg/g. It gradually declined over time, reaching 17.4 µg/g by the 4-hour time point. A similar decreasing trend in lithium content was observed in blood (declining from 39 µg/g to 15 µg/g), skin (from 24.2 µg/g to 2 µg/g), liver (from 51 µg/g to 9.8 µg/g) and kidney (from 105.2 µg/g to 24.3 µg/g). Lithium concentrations in the brain following intraperitoneal administration of lithium chloride were slightly higher than following oral administration and was 9.5–10.6 µg/g. The tumor/blood lithium concentration ratio (T/B) remained at 1.4-1.5 during the first three hours post-administration and reached 1.1 at 4 hours. The tumor/skin concentration ratio (T/S) exceeded 3 at all time points, specifically measuring 4.7 at the 1-hour time point.

**Table 2.** The lithium concentrations in tissues of the B16 melanoma-bearing mice after intraperitoneal lithium chloride administration (M±SD)

| **Group<sup>1</sup>**<br>**Time<sup>2</sup>** / | **1h** | **2h** | **3h** | **4h** |
| --- | --- | --- | --- | --- |
| **Tumor** | 54.1 ± 5.7 | 35.7 ± 9.9 | 25.1 ± 7.8 | 17.4 ± 8.2 |
| **Blood** | 39 ± 6.6 | 23.8 ± 6.6 | 18.1 ± 6.6 | 15 ± 3.9 |
| **Muscle** | 33.1 ± 12 | 19.4 ± 12.1 | 24.1 ± 12.9 | 21.2 ± 11.6 |
| **Skin** | 24.2 ± 14.8 | 9.4 ± 5.3 | 4.7 ± 4.8 | 2 ± 3.9 |
| **Liver** | 51 ± 8.9 | 23.4 ± 9.6 | 14.4 ± 4.9 | 9.8 ± 3.1 |
| **Kidney** | 105.2 ± 14.7 | 56.1 ± 13.7 | 34.9 ± 11.8 | 24.3 ± 7.5 |
| **Brain** | 10.6 ± 1.6 | 10.4 ± 3.4 | 10.3 ± 4.9 | 9.5 ± 4 |
| **T/S<sup>3</sup>** | 4.7 ± 5.4 | 3.5 ± 0.6 | 13.7 ± 18.9 | 6.7 ± 6.2 |
| **T/B<sup>4</sup>** | 1.4 ± 0.3 | 1.5 ± 0.3 | 1.4 ± 0.2 | 1.1 ± 0.4 |

<sup>1</sup>The number of animals was 6 in “1h” and “2h” groups, and 5 – in “3h” and “4h” groups.
<sup>2</sup>Time after lithium compound administration (hours).
<sup>3</sup>T/S: Tumor to Skin ratio.
<sup>4</sup>T/B: Tumor to Blood ratio.

A comparative analysis on the influence of the two lithium chloride administration routes on lithium biodistribution was carried out. The highest lithium concentration in the tumor was detected 1 hour after intraperitoneal administration, measuring 54.1 µg/g. At this time point, the tumor/skin lithium concentration ratio and the tumor/blood lithium concentration ratio were 4.7 and 1.4, respectively, representing the most favorable values compared to other time points. The highest tumor lithium concentration after oral administration was 30.2 µg/g, observed 4 hours post-administration. The most favorable tumor/skin and tumor/blood ratios were also found at the 4 hour time point, measuring 4.1 and 1.5, respectively. Thus, the optimal protocol to initiate neutron capture therapy is one hour after intraperitoneal administration.

Figure 2 shows the animal weights, Kaplan-Meier survival curves, and tumor growth dynamics for each group after intraperitoneal lithium administration. The mice in the LiNCT group weighed less than the three control groups, which, as in the model with oral lithium administration, could be due to the smaller tumor size in LiNCT-treated animals. The log-rank test showed a significant increase in survival in the LiNCT group compared to the other three control groups (Table 3). Moreover, the LiNCT group had statistically significant slower tumor growth (2-4 times) compared to the control groups in the first three weeks of the experiment (Table 4). Thus, LiNCT demonstrated a pronounced cytotoxic effect on tumor cells at the initial stages after irradiation.

**Table 3.** The LiNCT effects on survival rates of B16-bearing mice after intraperitoneal lithium administration.

| **Group** | **N (animals per group)** | **Mean ± SD** | **Median** | **ILS (%)<sup>1</sup>** | **P-value<sup>2</sup>** |
| --- | --- | --- | --- | --- | --- |
| Control | 11 | 14.0 ± 5.2 | 15 | 0 | - |
| Li | 12 | 11.1 ± 4.3 | 10 | -33.3 | 0.11815 vs Control |
| Neu | 12 | 19.8 ± 3.9 | 20 | 33.3 | 0.01688 vs Control<br>0.00023 vs Li |
| LiNCT | 10 | 33.6 ± 7.3 | 34 | 127 | 0.000017 vs Control<br>0.000015 vs Li<br>0.00019 vs Neu |

<sup>1</sup>Increase in life span (%) was calculated relative to the mean survival time of the Control group.
<sup>2</sup>The log-rank test with Holm's correction for multiple comparisons was used to estimate p-values based on the survival analysis results of the experimental animals.

**Table 4.** Statistically significant estimated effects of three experimental groups (Control, Li, Neu) relative to the LiNCT reference group on tumor volume dynamics at different time points (days), derived from a generalized linear mixed-effects model (GLMM).

| **Group<sup>1</sup>** | **Estimate<sup>2</sup>** | **Z-value<sup>3</sup>** | **P-value** |
| --- | --- | --- | --- |
| **Control\*3d** | 0.459 | 2.126 | 0.033 |
| **Li\*3d** | 0.944 | 4.380 | < 0.000001 |
| **Control\*7d** | 1.467 | 6.602 | < 0.000001 |
| **Li\*7d** | 2.047 | 9.493 | < 0.000001 |
| **Neu\*7d** | 0.731 | 3.457 | 0.001 |
| **Control\*9d** | 1.584 | 7.126 | < 0.000001 |
| **Li\*9d** | 1.838 | 8.261 | < 0.000001 |
| **Neu\*9d** | 0.905 | 4.279 | < 0.000001 |
| **Control\*11d** | 1.485 | 6.419 | < 0.000001 |
| **Li\*11d** | 1.656 | 6.632 | < 0.000001 |
| **Neu\*11d** | 0.665 | 3.153 | 0.002 |
| **Control\*14d** | 1.685 | 7.041 | < 0.000001 |
| **Li\*14d** | 2.055 | 7.193 | < 0.000001 |
| **Neu\*14d** | 0.924 | 4.319 | < 0.000001 |
| **Control\*16d** | 1.745 | 6.172 | < 0.000001 |
| **Li\*16d** | 2.168 | 6.739 | < 0.000001 |
| **Neu\*16d** | 0.931 | 4.293 | < 0.000001 |
| **Control\*18d** | 1.709 | 6.019 | < 0.000001 |
| **Neu\*18d** | 1.225 | 5.313 | < 0.000001 |
| **Control\*21d** | 1.475 | 3.578 | < 0.000001 |
| **Neu\*21d** | 1.111 | 4.502 | < 0.000001 |
| **Neu\*23d** | 0.780 | 3.012 | 0.003 |

<sup>1</sup>Interaction term between Group and Time factors.
<sup>2</sup>The logarithm of the ratio of the mean tumor volume for an indicated group and day compared to the LiNCT reference group on the same day.
<sup>3</sup>Z-statistic from the test of the estimated effect.

**Figure 2.**
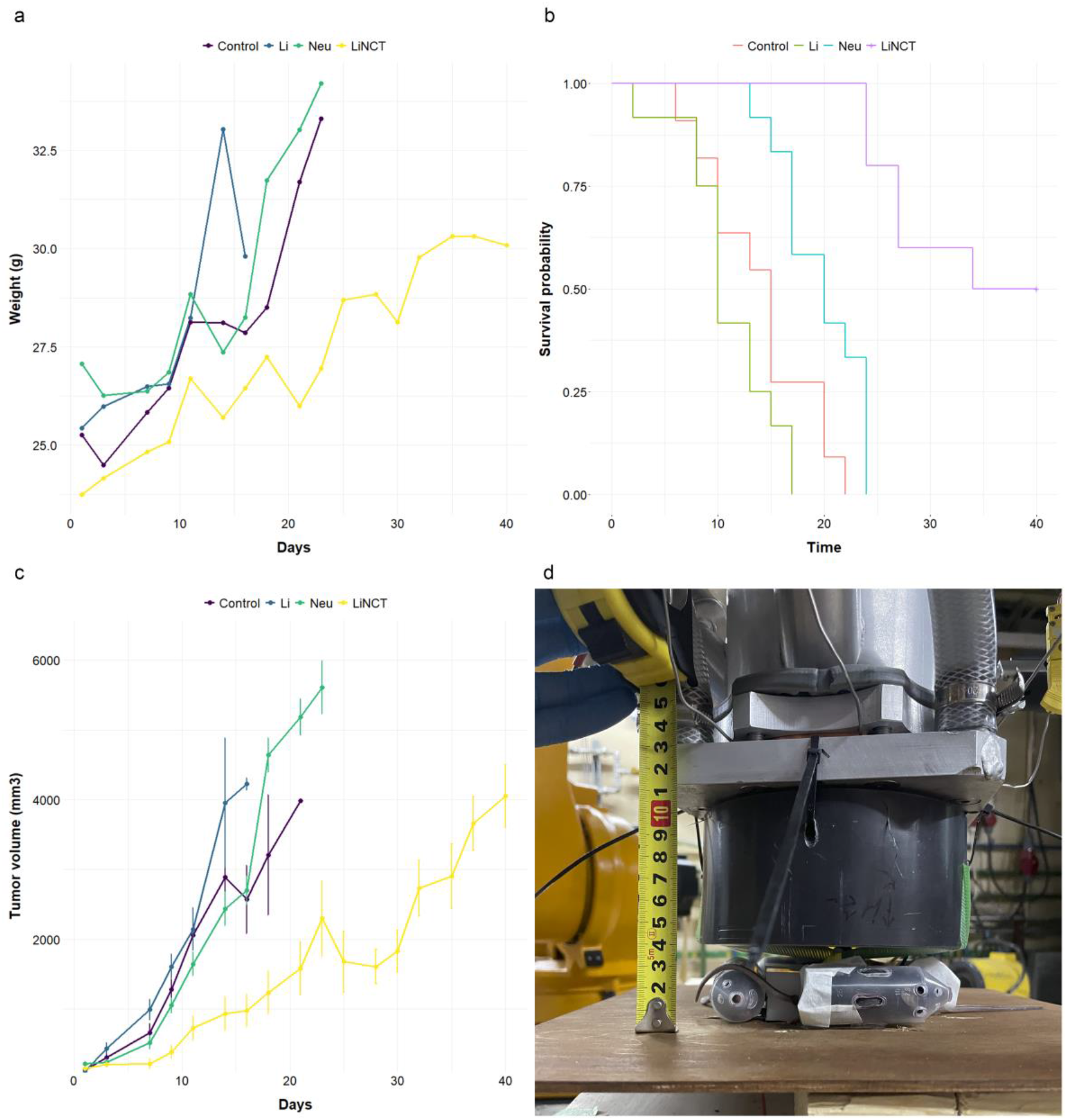
Evaluation of survival rate, animal weight and tumor growth dynamics for each experimental group after intraperitoneal lithium administration. (**a**) This figure shows the results of the animal weight dynamics (mean values) for B16 skin melanoma model after LiNCT and in control groups. (**b**) Survival times (days) are plotted, statistical analysis was performed using the log-rank test with Holm’s correction for multiple comparisons (“Control”: n = 11, “Li”: n = 12, “Neu”: n = 12, “LiNCT”: n = 10). (**c**) Tumor volume dynamics was analyzed using a generalized linear mixed-effects model (GLMM), the data are presented as means ± SD. (**d**) The representative image of experimental animals positioned in a specially designed restrictor under a collimator for neutron irradiation.

## Discussion

Our study provides the first comprehensive preclinical evaluation of LiNCT using peroral and intraperitoneal administration of a lithium compound in a murine tumor model. First of all, it is necessary to note what we have identified route-dependent tumor uptake of lithium. Intraperitoneal lithium administration achieved a significantly higher peak tumor lithium concentration (54 μg/g at 1h) compared to peroral drug administration (28-30 μg/g). This aligns with the established principle that intraperitoneal drug delivery often provides faster systemic absorption and higher bioavailability than the peroral route for many compounds.

According to our preliminary theoretical calculations, at the accumulation of 40 μg/g of lithium-6 in tumor cells the contribution of the dose of the nuclear reaction of neutron absorption by lithium will be 67% of the weighted total dose and 55% of the total physical dose, which is sufficient for therapy with selective accumulation of lithium in tumor cells^22^. It is known that for BNCT, the boron concentration in the tumor should be at least 15–30 μg/g of ^10^B atoms per gram of tumor tissue^25^, however, a number of studies have shown the success of BNCT at relatively low boron concentrations in the tumor, as well as high heterogeneity of boron accumulation^26-28^. In this *in vivo* study, the assessment of lithium biodistribution showed that lithium concentrations in the tumor were sufficient for LiNCT within one hour after both routes of lithium chloride administration and gradually decreased over time. The lithium compound we used (lithium chloride) was a non-selective lithium delivery agent, and we can assume that a more selective lithium-containing agent will be able to improve the targeting of lithium delivery to tumor cells. The tumor/blood ratio for animals exposed to peroral lithium compound administration was in the range of 0.8–1.5, tumor to skin ratio was in the range of 1.5–4.1, and blood to skin ratio was in the range of 1.5–3. Intraperitoneal lithium administration led to the next ranges ratios: 1.1-1.5 (tumor/blood), and 3.5-13.7 (tumor/skin).

It should be noted that fairly high values of lithium concentrations were obtained in the kidneys at all time points and with two routes of drug administration (oral and intraperitoneal). It is known that lithium is excreted primarily by the kidneys, as is boron, which explains the high concentrations of these elements in the kidneys^27,29^. Our previous series of experiments showed that administration of lithium carbonate at a dose of 400 mg/kg (corresponding to the dose we used in the experiment – 448.4 mg/kg lithium chloride) did not lead to significant nephrotoxicity, despite high lithium concentrations in the kidneys. Thus, we can assume that a single administration of lithium chloride at a dose of 450 mg/kg may be quite safe^22,24^.

A series of *in vivo* neutron irradiation experiments using the B16 skin melanoma model clearly demonstrated the efficacy of LiNCT. Higher lithium concentrations in the tumor with intraperitoneal administration, compared to the oral route, contributed to a significant increase in animal survival and a significant decrease in tumor growth compared to the three control groups.

The discussion of the dose received from irradiation deserves special attention due to the complexity. Recall that in BNCT, the total absorbed dose is the sum of four dose components: boron dose; nitrogen dose; fast neutron dose; γ-ray dose. A decade ago, it was believed that *“The first two dose components cannot be measured in principle”*, as was previously written here^10^ (p. 279). The methods for measuring the fast neutron dose for BNCT are absent also, as the energy of neutrons is obviously lower than 1 MeV and, for example, fission ionization chambers are not applicable. There are quite a few proven approaches for measuring of γ-ray dose alone. When evaluating the patient’s response to the therapy, it is customary to analyze the equivalent (biological) dose, since the destructive effect of different ionizing particles can be different. The damage of tissue is due to three types of directly ionizing radiations that differ in their linear energy transfer (LET) characteristics: low-LET γ-rays, high-LET protons and high-LET heavier charged particles ^4^He, ^7^Li, and ^14^C. The equivalent dose is the sum of four dose components each of which is multiplied by a weighted factor. Some of these weighted factors are not well defined, and the weighted factor of boron doze depends on the boron carrier used and the tissue. All this demonstrates how dosimetry in BNCT is more complicated than in photon and electron therapy, which both act by energy transfer via electrons. Although significant progress has been made in the development of dosimetry tools and methods over the past decade, and some of them have been developed by us and are listed in the Methods chapter, dosimetry in BNCT remains a complex issue largely due to the need to know the concentration of boron in tumor cells and in cells of healthy organs. The hope is to use the prompt *γ*-ray spectroscopy, since the nuclear reaction of neutron absorption by boron results in the emission of 478 keV photon. Also, for a preliminary assessment of boron accumulation, boron-phenylalanine labeled with fluorine-18 is used, the decay of which is recorded by a PET tomography.

In LiNCT, things are even more difficult, since the method of prompt *γ*-ray spectroscopy is not applicable (the reaction products are only charged particles) and there is no lithium drug labeled with a radionuclide for PET tomography. For this reason, when choosing the irradiation regimen, we relied on our ten years of experience working with boron, which we gained through irradiating cell cultures and laboratory animals^30-36^, testing new boron delivery drugs^37-41^, and treating domestic cats and dogs with spontaneous tumors^42,43^.

Finally, two important circumstances must be kept in mind also. Firstly, the movement of a neutron in the human body (water phantom) is fundamentally different from the almost rectilinear propagation of radiation in photon, proton, heavy-ion therapy. A neutron, scattering on the atomic nuclei and slowing down, moves chaotically like a Brownian particle. It is impossible to predict where a neutron will be captured by boron or lithium, we can only say that in an infinite medium a neutron is captured by boron or lithium in a volume of about 1 liter. That is, even if you made a pencil beam of neutrons, nuclear reactions will occur in a larger volume than the size of the beam. For this reason, laboratory mice are not a very convenient object for demonstrating the effectiveness of neutron capture therapy. Not only the tumor is exposed to radiation, but also the kidneys and liver, in which boron (lithium) has accumulated in an even higher concentration than in the tumor. Therefore, it is necessary to limit the radiation dose so as not to kill the mouse, damaging the kidneys and liver. It is very difficult, sometimes almost impossible, to ensure such a duration of irradiation that, on the one hand, tumors are completely destroyed, and at the same time, on the other hand, other vital organs are not damaged, so that the mouse continues to live during the entire observation period. Secondly since the cross-section of the ^6^Li(n,α)^3^H reaction is four times smaller than the cross section of the ^10^B(n,α)^7^Be reaction, and the energy released in the reaction, on the contrary, is twice as large, then for the same effect, twice as much lithium is required as boron.

## Conclusions

The results of our study provide completely new opportunities for the development of neutron capture therapy: both with lithium and with a possible lithium-boron combination. The concept of NCT, assuming selectivity of the neutron capture agent accumulation and high linear energy transfer, may indeed provide significant efficiency of this method in comparison with other types of radiotherapy, thus we strongly encourage researchers to conduct further studies to confirm the clinical applicability of LiNCT.

## Materials and Methods

### Lithium compound

Lithium chloride with a lithium-6 mass fraction of more than 90 % (LiCl) was purchased from the Novosibirsk Chemical Concentrates Plant (Novosibirsk, Russia), an enterprise of the State Atomic Energy Corporation Rosatom.

### Experimental design using B16 melanoma model in vivo

Male C57BL/6 mice of 10-12 weeks of age, with weights of 20-22 g were obtained from the Institute of Cytology and Genetics, Siberian Branch, Russian Academy of Sciences, Novosibirsk. All the animals were housed in an environment with constant room temperature (23 °C), natural day/night light cycle, and standard laboratory food and water were provided. All experiments were performed in accordance with the EU Directive 2010/63/EU and the ARRIVE guidelines. The study protocol was approved by the Ethics Committee of the Research Institute of Clinical and Experimental Lymphology (Approval No. 194, 29 November 2024).

Two models were used in the experiments: the B16 tumor model with peroral administration of lithium chloride enriched with ^6^Li and the B16 tumor model with intraperitoneal administration of lithium chloride enriched with ^6^Li at a dose of 450 mg/kg. The series of experiments was divided into 4 stages and was carried out using a total of 115 mice with implanted skin melanoma. *Study 1* included of 20 animals and consisted in assessing the lithium accumulation in the organs of mice after oral drug administration at 4 time points (1, 2, 3, and 4 hours). Then, LiNCT was performed on the B16 tumor model with oral administration of lithium chloride (*study 2*, 28 animals). After analyzing the results, we decided to assess lithium accumulation in 22 mice administered lithium chloride intraperitoneally (*study 3*) at the same time points as in study 1 (1, 2, 3, and 4 hours). Finally, LiNCT was performed on the B16 tumor model with intraperitoneal administration of the drug (*study 4*, 45 animals).

For tumor induction, cultured B16 cells were injected subcutaneously into the right inguinal region of mice (2 × 10^6^ cells). After tumor growth induction (10 days in the *per os* model and 7 days in the intraperitoneal drug administration model), mice were randomly divided into groups for each experiment: to assess lithium biodistribution – 1, 2, 3, and 4 hours; to assess the efficacy of LiNCT - into “Control” (intact tumor), “Li” (control group, obtained lithium compound only), “Neu” (neutron irradiation control group), and “LiNCT” (neutron irradiation 1-1.5 h after lithium compound administration) groups. Animals were observed until death or euthanasia by cervical dislocation (without anesthesia) in compliance with the approved study protocol. The efficacy of LiNCT was assessed by tumor growth dynamics and survival of experimental animals. Animal weights and tumor volume were measured every 3-4 days. The tumor volume was calculated by the formula: V = (Width^2^ × Length) / 2. The percentage increase in life span (ILS) was calculated by the formula: (median survival time of experimental group – median survival time of control group) × 100 / (median survival time of control group)^44^.

### Lithium biodistribution in vivo in B16 melanoma model

The inductively coupled plasma atomic emission spectrometry (ICP AES) on an ICPE-9820 high-resolution spectrometer (Shimadzu, Japan) was used to measure lithium concentrations in tumor and organs of interest. Lithium chloride was administered at a dose of 450 mg/kg: orally at a volume of 20 μL per animal and intraperitoneally at a volume of 400 μL per animal. Euthanasia was performed 1, 2, 3, and 4 hours after lithium chloride administration. Tumor, blood, muscle, skin, liver, kidney and brain were excised and weighed. The normal tissue samples were obtained at a distance of 1-2 mm from the tumor surgical margin. Tissue sample preparation was performed using concentrated HNO_3_ (high-purity grade, 69%) and hydrogen peroxide H_2_O_2_ (37%). Samples were heated to 90°C in a Dry Block Heater 2 (IKA, Germany) until complete dissolution and formation of a clear solution. Each sample volume was adjusted to 6 mL with deionized water. Samples were introduced via a peristaltic pump. Calibration curves were constructed using a certified single-element lithium ion reference material Lithium (Li) CRISTAR® Single Element Std. Soln. for ICP (CDH Ltd., India). Analytical results were obtained by 670.784 emission line. Final lithium concentrations in tumors and organs were calculated using the formula: measured concentration × sample volume / organ weight.

### Neutron irradiation

Animals were irradiated using the accelerator based neutron source VITA at the Budker Institute of Nuclear Physics in Novosibirsk, Russia^45-47^. Neutrons were generated in ^7^Li(p,n)^7^Be reaction at a proton energy of 2.1 MeV a current of 1.6 mA for about 1.5 hours, gaining a fluence of 2.5 mA·h. The lithium target for generating neutrons is placed in the vertical tract of the facility. A 155 mm diameter, 72 mm high polyethylene disk with a volumetric bismuth inclusion is placed tightly below the target. This disk serves as a moderator to slow the neutrons to thermal energies so that they are effectively captured by lithium-6, leading to the nuclear reaction ^6^Li(n,α)^3^H with a large energy release.

Below the moderator in specially designed restrictors, the animals are placed, placing the leg of the animal with the tumor towards the center and closer to the center, and the body on the periphery. For peroral administration, up to 6 animals are placed this way, for intraperitoneal administration – 4. For intraperitoneal administration, plexiglass plates are also placed between the legs of neighboring animals in order to slightly increase the dose to the tumor. A 20 mm high moderator-like disk is placed under the animals to ensure more uniform irradiation with thermal neutrons due to their scattering. All this is placed on a platform held by a robotic arm. By controlling the robotic arm, the animals are delivered to the irradiation zone and removed from the irradiation zone after irradiation.

### Dosimetry

To measure the dose of ionizing radiation in a radiation-protected bunker where the VITA is located and in the control room, a stationary installed DBG-S11D γ-radiation dosimeter, BDMN-100-07 neutron detection device and BOP-1M data processing and transmission unit («Doza», Russia), LB6500-3H-10 γ-detector with a Micro Gamma LB 112 display unit (Berthold Technologies, Germany) and a DKS-96 portable dosimeter-radiometer with BDMN-96 and BDMG-96 detection units («Doza», Russia) are used.

A small-sized detector with a pair of cast polystyrene scintillators, one of which is enriched with boron, is used to measure the boron dose and the *γ*-ray dose in the air and in a water phantom (it should be noted that the lithium dose is half the boron dose). A lithium-containing scintillator GS20 (The Saint-Gobain Crystals, USA) mounted on a Hamamatsu R6095 photomultiplier with a high-voltage power source MHV12-1.5K1300P (TRACO Electronics, Japan) is used to register neutrons. The proposed and developed “cellular dosimeter” is used to estimate the contribution of the sum of doses of fast neutrons and thermal neutrons^34^. A SEG-1KP *γ*-radiation spectrometer based on a semiconductor detector made of high-purity germanium (Institute of Physical and Technical Problems, Dubna, Russia) is used to measure the boron dose by the prompt *γ*-ray spectrometry method during therapy^48^.

### Statistical analysis

All data were analyzed by using R language (version 4.4.2) [R Foundation for Statistical Computing R: A Language and Environment for Statistical Computing. R Core Team. http://www.r-project.org Accessed on 28 July 2025]. The normality of the data was evaluated using the Shapiro-Wilk test, with a significance criterion of p<0.05. Tumor volume dynamics was analyzed using a generalized linear mixed-effects model (GLMM) with a Gamma distribution and log link function to appropriately model the positive, right-skewed nature of tumor volume data. The model included fixed effects for experimental Group, Time (days), and their interaction to evaluate differences in tumor growth dynamics among groups. Statistical analyses were performed using the glmmTMB package^49^. Kaplan–Meier curves were used to evaluate survival times. The log-rank test with Holm’s correction for multiple comparisons was used to estimate p-values based on the survival analysis results of the experimental animals. Two-tailed p < 0.05 deemed statistical significance.

## Author Contributions

Conceptualization: I. Taskaeva and S. Taskaev; Data analysis: I. Taskaeva, A. Kasatova, T. Bykov, V. Richter, E. Sokolova, N. Bgatova and S. Taskaev; Data interpretation: I. Taskaeva, A. Kasatova, N. Bgatova and S. Taskaev; Write manuscript: I. Taskaeva, A. Kasatova and S. Taskaev.

## Availability of materials and data

The datasets generated during and/or analysed during the current study are available from the corresponding author on reasonable request.

## References

1. Suzuki, M. Boron neutron capture therapy (BNCT): a unique role in radiotherapy with a view to entering the accelerator-based BNCT era. Int. J. Clin. Oncol. 25, 43–50 (2020).

2. Ahmed, M. et al. Advances in boron neutron capture therapy. International Atomic Energy Agency. Vienna, Austria. p. 416. (2023).

3. Matsumoto, Y., Fukumitsu, N., Ishikawa, H., Nakai, K. & Sakurai, H. A critical review of radiation therapy: from particle beam therapy (proton, carbon, and BNCT) to beyond. J. Pers. Med. 11, 825 (2021).

4. Locher, G. L. Biological effects and therapeutic possibilities of neutrons. Am. J. Roentgenol. Radium Ther. Nucl. Med. 36, 1–13 (1936).

5. Jin, W. H., Seldon, C., Butkus, M., Sauerwein, W. & Giap, H. B. A review of boron neutron capture therapy: its history and current challenges. Int. J. Part. Ther. 9, 71–82 (2022).

6. Cheng, X., Li, F. & Liang, L. Boron neutron capture therapy: clinical application and research progress. Curr. Oncol. 29, 7868–7886 (2022).

7. Sauerwein, W. A. G., Sancey, L., Hey-Hawkins, E., Kellert, M., Panza, L., Imperio, D., Balcerzyk, M., Rizzo, G., Scalco, E., Herrmann, K., Mauri, P., De Palma, A. & Wittig, A. Theranostics in boron neutron capture therapy. Life (Basel) 11, 330 (2021).

8. Barth, R. F., Mi, P. & Yang, W. Boron delivery agents for neutron capture therapy of cancer. Cancer Commun. (Lond.) 38, 35 (2018).

9. Dymova, M. A., Taskaev, S. Y., Richter, V. A. & Kuligina, E. V. Boron neutron capture therapy: current status and future perspectives. Cancer Commun. (Lond.) 40, 406–421 (2020).

10. Sauerwein, W. A. G., Wittig, A., Moss, R. & Nakagawa, Y. Neutron capture therapy: principles and applications. Springer, Berlin/Heidelberg, Germany (2012).

11. Cade, J. F. Lithium salts in the treatment of psychotic excitement. Med. J. Aust. 2, 349–352 (1949).

12. Mitchell, P. B. & Hadzi-Pavlovic, D. Lithium treatment for bipolar disorder. Bull. World Health Organ. 78, 515–517 (2000).

13. Malhi, G. S., Tanious, M., Das, P. & Berk, M. The science and practice of lithium therapy. Aust. N. Z. J. Psychiatry 46, 192–211 (2012).

14. Oruch, R., Elderbi, M. A., Khattab, H. A., Pryme, I. F. & Lund, A. Lithium: a review of pharmacology, clinical uses, and toxicity. Eur. J. Pharmacol. 740, 464–473 (2014).

15. Wen, J., Sawmiller, D., Wheeldon, B. & Tan, J. A review for lithium: pharmacokinetics, drug design, and toxicity. CNS Neurol. Disord. Drug Targets 18, 769–778 (2019).

16. Kakhki, S. & Ahmadi-Soleimani, S. M. Experimental data on lithium salts: from neuroprotection to multi-organ complications. Life Sci. 306, 120811 (2022).

17. Zahl, P. A. & Cooper, F. S. Localization of lithium in tumor tissue as a basis for slow neutron therapy. Science 93, 64–65 (1941).

18. Gorkin, R. A. & Richelson, E. Lithium ion accumulation by cultured glioma cells. Brain Res. 171, 365–368 (1979).

19. Gorkin, R. A. & Richelson, E. Lithium transport by mouse neuroblastoma cells. Neuropharmacology 20, 791–801 (1981).

20. Rendina, L. M. Can lithium salts herald a new era for neutron capture therapy? J. Med. Chem. 53, 8224–8227 (2010).

21. Gonçalves, G., Sandoval, S., Llenas, M., Ballesteros, B., Da Ros, T., Bortolussi, S., Cansolino, L., Ferrari, C., Postuma, I., Protti, N., Melle-Franco, M., Altieri, S. & Tobías-Rossell, G. Lithium halide filled carbon nanocapsules: paving the way towards lithium neutron capture therapy (LiNCT). Carbon 208, 148–159 (2023).

22. Taskaeva, I., Kasatova, A., Surodin, D., Bgatova, N. & Taskaev, S. Study of lithium biodistribution and nephrotoxicity in skin melanoma mice model: the first step towards implementing lithium neutron capture therapy. Life (Basel) 13, 518 (2023).

23. Taskaeva, I., Kasatova, A., Razumov, I., Bgatova, N. & Taskaev, S. Lithium salts cytotoxicity and accumulation in melanoma cells in vitro. J. Appl. Toxicol. 44, 712–719 (2024).

24. Taskaeva, Y. S., Kasatova, A. I., Shatruk, A. Y., Taskaev, S. Y. & Bgatova, N. P. The expression of markers of acute kidney injury Kim1 and NGAL after administration of high doses of lithium carbonate in mice with engrafted skin melanoma B16. Bull. Exp. Biol. Med. 176, 567–571 (2024).

25. Sumitani, S., Oishi, M., Yaguchi, T., Murotani, H., Horiguchi, Y., Suzuki, M., Ono, K., Yanagie, H. & Nagasaki, Y. Pharmacokinetics of core-polymerized, boron-conjugated micelles designed for boron neutron capture therapy for cancer. Biomaterials 33, 3568–3577 (2012).

26. Garabalino, M. A., Heber, E. M., Monti Hughes, A., González, S. J., Molinari, A. J., Pozzi, E. C., Nievas, S., Itoiz, M. E., Aromando, R. F., Nigg, D. W., Bauer, W., Trivillin, V. A. & Schwint, A. E. Biodistribution of sodium borocaptate (BSH) for boron neutron capture therapy (BNCT) in an oral cancer model. Radiat. Environ. Biophys. 52, 351–361 (2013).

27. Carpano, M., Perona, M., Rodriguez, C., Nievas, S., Olivera, M., Santa Cruz, G. A., Brandizzi, D., Cabrini, R., Pisarev, M., Juvenal, G. J. & Dagrosa, M. A. Experimental studies of boronophenylalanine ((10)BPA) biodistribution for the individual application of boron neutron capture therapy (BNCT) for malignant melanoma treatment. Int. J. Radiat. Oncol. Biol. Phys. 93, 344–352 (2015).

28. Zhang, Z., Yong, Z., Jin, C., Song, Z., Zhu, S., Liu, T., Chen, Y., Chong, Y., Chen, X. & Zhou, Y. Biodistribution studies of boronophenylalanine in different types of skin melanoma. Appl. Radiat. Isot. 163, 109215 (2020).

29. Timmer, R. T. & Sands, J. M. Lithium intoxication. J. Am. Soc. Nephrol. 10, 666–674 (1999).

30. Mostovich, L. A., Gubanova, N. V., Kutsenko, O. S., Aleinik, V. I., Kuznetsov, A. S., Makarov, A. N., Sorokin, I. N., Taskaev, S. Y., Nepomnyashchikh, G. I. & Grigor’eva, E. V. Effect of epithermal neutrons on viability of glioblastoma tumor cells in vitro. Bull. Exp. Biol. Med. 151, 264–267 (2011).

31. Sato, E., Zaboronok, A., Yamamoto, T., Nakai, K., Taskaev, S., Volkova, O., Mechetina, L., Taranin, A., Kanygin, V., Isobe, T., Mathis, B. J. & Matsumura, A. Radiobiological response of U251MG, CHO-K1 and V79 cell lines to accelerator-based boron neutron capture therapy. J. Radiat. Res. 59, 101–107 (2018).

32. Taskaev, S. Development of an Accelerator-Based Epithermal Neutron Source for Boron Neutron Capture Therapy. Phys. Part. Nucl. 50, 569–575 (2019).

33. Zavjalov, E., Zaboronok, A., Kanygin, V., Kasatova, A., Kichigin, A., Mukhamadiyarov, R., Razumov, I., Sycheva, T., Mathis, B. J., Maezono, S. E. B., Matsumura, A. & Taskaev, S. Accelerator-based boron neutron capture therapy for malignant glioma: a pilot neutron irradiation study using boron phenylalanine, sodium borocaptate and liposomal borocaptate with a heterotopic U87 glioblastoma model in SCID mice. Int. J. Radiat. Biol. 96, 868–878 (2020).

34. Dymova, M., Dmitrieva, M., Kuligina, E., Richter, V., Savinov, S., Shchudlo, I., Sycheva, T., Taskaeva, I. & Taskaev, S. Method of measuring high-LET particles dose. Radiat. Res. 196, 192–196 (2021).

35. Kanygin, V. V., Kasatova, A. I., Zavjalov, E. L., Razumov, I. A., Kolesnikov, S. I., Kichigin, A. I., Solov’eva, O. I., Tsygankova, A. R., Taskaev, S. Y., Kasatov, D. A., Sycheva, T. V. & Byvaltsev, V. A. Effects of boron neutron capture therapy on the growth of subcutaneous xenografts of human colorectal adenocarcinoma SW-620 in immunodeficient mice. Bull. Exp. Biol. Med. 172, 356–361 (2021).

36. Kanygin, V., Razumov, I., Zaboronok, A., Zavjalov, E., Kichigin, A., Solovieva, O., Tsygankova, A., Guselnikova, T., Kasatov, D., Sycheva, T., Mathis, B. J. & Taskaev, S. Dose-dependent suppression of human glioblastoma xenograft growth by accelerator-based boron neutron capture therapy with simultaneous use of two boron-containing compounds. Biology 10, 1124 (2021).

37. Vorobyeva, M. A., Dymova, M. A., Novopashina, D. S., Kuligina, E. V., Timoshenko, V. V., Kolesnikov, I. A., Taskaev, S. Y., Richter, V. A. & Venyaminova, A. G. Tumor Cell-Specific 2’-Fluoro RNA Aptamer Conjugated with Closo-Dodecaborate as a Potential Agent for Boron Neutron Capture Therapy. Int. J. Mol. Sci. 22, 7326 (2021).

38. Popova, T., Dymova, M. A., Koroleva, L. S., Zakharova, O. D., Lisitskiy, V. A., Raskolupova, V. I., Sycheva, T., Taskaev, S. & Silnikov, V. N., Godovikova, T. S. Homocystamide conjugates of human serum albumin as a platform to prepare bimodal multidrug delivery systems for boron-neutron capture therapy. Molecules 26, 6537 (2021).

39. Zaboronok, A., Khaptakhanova, P., Uspenskii, S., Bekarevich, R., Mechetina, L., Volkova, O., Mathis, B. J., Kanygin, V., Ishikawa, E., Kasatova, A., Kasatov, D., Shchudlo, I., Sycheva, T., Taskaev, S. & Matsumura, A. Polymer-Stabilized Elemental Boron Nanoparticles for Boron Neutron Capture Therapy: Initial Irradiation Experiments. Pharmaceutics 14, 761 (2022).

40. Aiyyzhy, K. O., Barmina, E. V., Zavestovskaya, I. N., Kasatova, A. I., Petrunya, D. S., Uvarov, O. V., Saraykin, V. V., Zhilnikova, M. I., Voronov, V. V. & Shafeev, G. A. Laser ablation of Fe2B target enriched in 10sB content for boron neutron capture therapy. Laser Phys. Lett. 19, 066002 (2022).

41. Zavestovskaya, I. N., Kasatova, A. I., Kasatov, D. A., Babkova, J. S., Zelepukin, I. V., Kuzmina, K. S., Tikhonowski, G. V., Pastukhov, A. I., Aiyyzhy, K. O., Barmina, E. V., Popov, A. A., Razumov, I. A., Zavjalov, E. L., Grigoryeva, M. S., Klimentov, S. M., Ryabov, V. A., Deyev, S. M., Taskaev, S. Y. & Kabashin, A. V. Laser-synthesized elemental boron nanoparticles for efficient boron neutron capture therapy. Int. J. Mol. Sci. 24, 17088 (2023).

42. Kanygin, V., Kichigin, A., Zaboronok, A., Kasatova, A., Petrova, E., Tsygankova, A., Zavjalov, E., Mathis, B. J. & Taskaev, S. In vivo Accelerator-based Boron Neutron Capture Therapy for Spontaneous Tumors in Large Animals: Case Series. Biology 11, 138 (2022).

43. Kanygin, V., Zaboronok, A., Kichigin, A., Petrova, E., Guselnikova, T., Kozlov, A., Lukichev, D., Mathis, B. J. & Taskaev, S. Gadolinium neutron capture therapy for cats and dogs with spontaneous tumors using Gd-DTPA. Veterinary Sciences 10, 274 (2023).

44. Kayama, R., Tsujino, K., Kawabata, S., Fujikawa, Y., Kashiwagi, H., Fukuo, Y., Hiramatsu, R., Takata, T., Tanaka, H., Suzuki, M., Hu, N., Miyatake, S. I., Takami, T. & Wanibuchi, M. Translational research of boron neutron capture therapy for spinal cord gliomas using rat model. Sci. Rep. 14, 8265 (2024).

45. Taskaev, S., Berendeev, E., Bikchurina, M., Bykov, T., Kasatov, D., Kolesnikov, I., Koshkarev, A., Makarov, A., Ostreinov, G., Porosev, V., Savinov, S., Shchudlo, I., Sokolova, E., Sorokin, I., Sycheva, T., Verkhovod, G. Neutron Source Based on Vacuum Insulated Tandem Accelerator and Lithium Target. Biology 10, 350 (2021).

46. Taskaev, S. Neutron Source VITA. in: Advances in boron neutron capture therapy. International Atomic Energy Agency. Vienna, Austria, pp. 255–260 (2023).

47. Taskaev, S. Accelerator based Neutron Source VITA. Fizmatlit, Moscow, Russia, 248 p. (2024).

48. Bikchurina, M., Bykov, T., Ibrahim, I., Kasatova, A., Kasatov, D., Kolesnikov, I., Konovalova, V., Kormushakov, T., Koshkarev, A., Kuznetsov, A., Porosev, V., Savinov, S., Shchudlo, I., Singatulina, N., Sokolova, E., Sycheva, T., Taskaeva, I., Verkhovod, G. & Taskaev, S. Dosimetry for Boron Neutron Capture Therapy Developed and Verified at the Accelerator based Neutron Source VITA. Front. Nucl. Eng. 2, 1266562 (2023).

49. Brooks, M. E., Kristensen, K., van Benthem, K. J., Magnusson, A., Berg, C. W., Nielsen, A., Skaug, H. J., Maechler, M. & Bolker, B. M. glmmTMB balances speed and flexibility among packages for zero-inflated generalized linear mixed modeling. The R Journal 9, 378–400 (2017).

